# The autism-associated Neuroligin-3 R451C mutation alters mucus density and the spatial distribution of bacteria in the mouse gastrointestinal tract

**DOI:** 10.1101/2022.06.27.497808

**Authors:** Madushani Herath, Joel C Bornstein, Elisa L Hill-Yardin, Ashley E Franks

## Abstract

The intestinal mucus layer protects the host from invading pathogens and is essential for maintaining a healthy mucosal microbial community. Alterations in the mucus layer and composition of mucus-residing microbiota in people diagnosed with Autism Spectrum Disorder (ASD; autism) may contribute to dysbiosis and gastrointestinal (GI) dysfunction. Although microbial dysbiosis based on sequencing data is frequently reported in autism, spatial profiling of microbes adjacent to the mucosa is needed to identify changes in bacterial subtypes in close contact with host tissues. Here, we analysed the spatial distribution of the MUC-2 protein using immunofluorescence as well as total bacteria, Bacteroidetes, Firmicutes phyla and *Akkermansia muciniphila* (*A. muciniphila*) using fluorescent *in situ* hybridization in mice expressing the autism-associated R451C mutation in the *Neuroligin-3 (Nlgn3)* gene. We show that the *Nlgn3* R451C mutation increases mucus density adjacent to the distal ileal epithelium in mice. The relative density of total bacteria, Firmicutes and *A. muciniphila* was increased whereas the density of Bacteroidetes was decreased closer to the epithelium in *Nlgn3*^*R451C*^ mice. In summary, this study suggests that increased mucus density could contribute to mucosal microbial dysbiosis in ASD.

## MAIN

The intestinal mucus layer demarcates the enteric microbiota from the interior of the body and serves as the sole source of energy for mucus-residing bacteria (Marcobal et al., 2013). Animal models with an impaired mucus layer are predisposed to gastrointestinal (GI) diseases, suggesting that mucus layer integrity is important for maintaining intestinal homeostasis (Van der Sluis et al., 2006, Lennon et al., 2014, Johansson et al., 2013). Patients with ASD are at high risk of hospitalization due to GI dysfunction (McElhanon et al., 2014) but the cause is currently unknown. Co-existence of GI dysfunction with an altered abundance of mucolytic bacteria in individuals with ASD (Wang et al., 2011, Luna et al., 2017) suggests that changes in the mucus composition could contribute to these individuals being prone to intestinal dysfunction.

Mice expressing the autism-associated R451C mutation encoding Neuroligin-3 (NLGN3), a postsynaptic membrane protein, show GI dysfunction (Hosie et al., 2019). Expression of NLGN3 in most enteric neurons and glial cells (Herath et al., 2022) suggests that NLGN3 plays an important role in neurally-mediated gut function including mucus secretion. Given that the enteric nervous system (ENS) contributes to maintaining the integrity of the mucus layer (Specian and Neutra, 1980) reviewed in (Herath et al., 2020) alterations in the enteric neural network potentially affect mucus properties. However, the impact of this mutation on intestinal mucus layer thickness and the distribution of microbial communities embedded in the mucus biofilm has not been investigated.

Here, we investigated the impact of the *Nlgn3* R451C mutation on mucus layer density and the mucosa-associated microbial community in the mouse GI tract. To accomplish this task, we combined a mucus preservation method (methanol Carnoy’s fixation) and fluorescent *in situ* hybridization to localize bacteria within the mucus layer of the mouse distal ileum. Dominant components of the GI microbiome in human and mice including the Bacteroidetes and

Firmicutes phyla were selected to characterise the spatial distribution of microbes adjacent to the mucosal surface in wild-type (WT) and *Nlgn3*^R451C^ mice. The ratio of abundance of these phyla is reportedly altered in individuals with ASD (Strati et al., 2017, Tomova et al., 2015). The spatial distribution of the mucus degrading species *A. muciniphila* was also analysed in the mouse ileum. The MATLAB based BacSpace image analysis platform (Earle et al., 2015) was used to evaluate the effects of the R451C mutation on mucus layer density variation and microbial localization patterns in the distal ileum. The relative mucus and bacterial density were determined by averaging the fluorescence signal intensity and normalising as a function of distance from the epithelium towards lumen (Earle et al., 2015).

We found that mucus layer density is altered in *Nlgn3*^R451C^ mice compared to WT (p**<**0.0001, n=5 in both groups). In both WT and *Nlgn3*^R451C^ mice, maximum mucus density occurred at approximately 11 μm from the mucus-epithelial border and was significantly higher in *Nlgn3*^R451C^ mutant mice compared to WT (WT: 0.84 ± 0.3, *Nlgn3*^R451C^ mutant mice: 2.55 ± 0.2 (normalized density), p=0.003, n=5; **Figure 1 A1-A4**).

**Figure 1.**
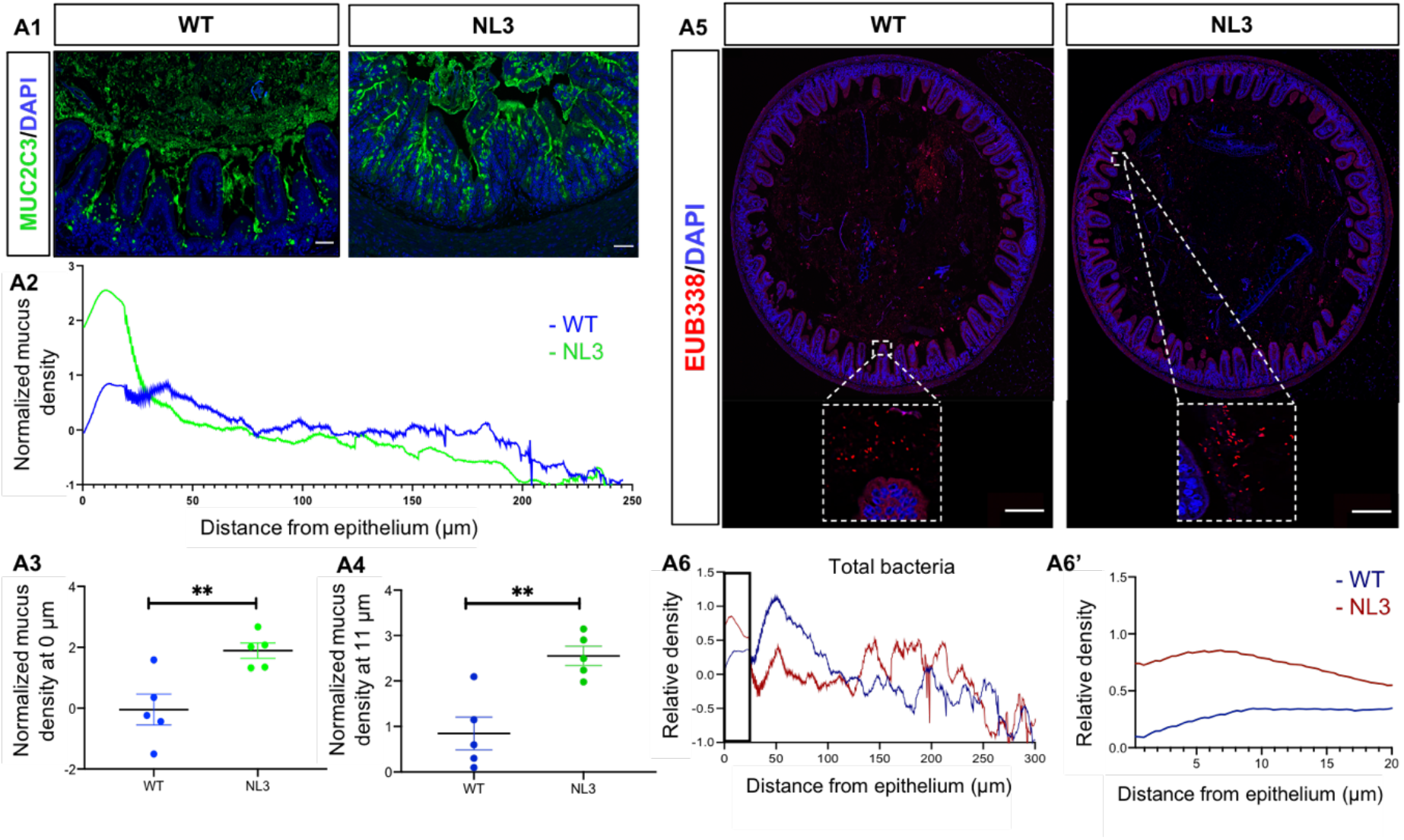
Ileal mucus layer density variation in the presence of the *Nlgn3* R451C mutation. Confocal images of transverse sections of the distal ileum in (**A1**) WT and *Nlgn3*^R451C^ mutant mice. **A2**: Mucus density variation from the mucosal to luminal direction averaged over the length of the epithelium in WT (blue) and *Nlgn3*^R451C^ mutant mice (green). **A3**: Comparison of the average mucus density at the epithelium in WT and *Nlgn3*^R451C^ mice. **A4**: Average mucus density comparison at 11 μm from the epithelium in WT and *Nlgn3*^R451C^ mice. **A5:** High-resolution confocal images of the distal ileum labelled with the universal bacterial marker EUB338, and DAPI, DNA marker in WT and *Nlgn3*^R451C^ mutant mice. White insets: bacterial density in close proximity to the epithelium. **A6**: Spatial distribution profile of total bacteria in WT and *Nlgn3*^R451C^ mutant mice. **A6’** (enlarged area of interest indicated by black rectangle in A6): local distribution pattern of bacteria within 20 μm from the epithelium. Data are mean-subtracted and divided by the standard deviation for normalization (n=5 in both groups). **P<0.01, scale bar=100 μm.

Since we observed clear differences in mucus density between genotypes within 0-20 μm from the mucus-epithelial border, we also analysed total bacterial density, Phylum Bacteroidetes, Phylum Firmicutes and *A. muciniphila* density in this region. Interestingly, the relative total bacterial density was increased in *Nlgn3*^R451C^ mutant mice compared to WT within this range (p<0.0001, n=5 in both groups) (**Figure A5 and A6**). *Nlgn3*^R451C^ mice showed a lower density of Bacteroidetes within the 0-20 μm region adjacent to the mucosa of the distal ileum compared to WT. These data show that the spatial pattern of Bacteroidetes is shifted by expression of the R451C mutation (p<0.0001, n=5 in both groups) (**Figure 2 B1-B2**). The distribution of Firmicutes within 20 μm of the epithelial boundary was also shifted such that the density of Firmicutes was higher in *Nlgn3*^R451C^ mice than in WT in ileal cross sections (WT; n=3, *Nlgn3*^R451C^ mutant mice; n=3, p<0.0001) (**Figure 2 B3-B4**). We show that *A. muciniphila* density is significantly reduced close to the mucosa (i.e., within 10 μm of the mucosa), but increased to above WT levels at a greater distance (i.e., further than 10 μm) from the mucosal epithelium in *Nlgn3*^R451C^ mutant mice (WT; n=4, *Nlgn3*^R451C^ mutant mice; n=5, p<0.0001) (**Figure 2 B5-B6**).

**Figure 2:**
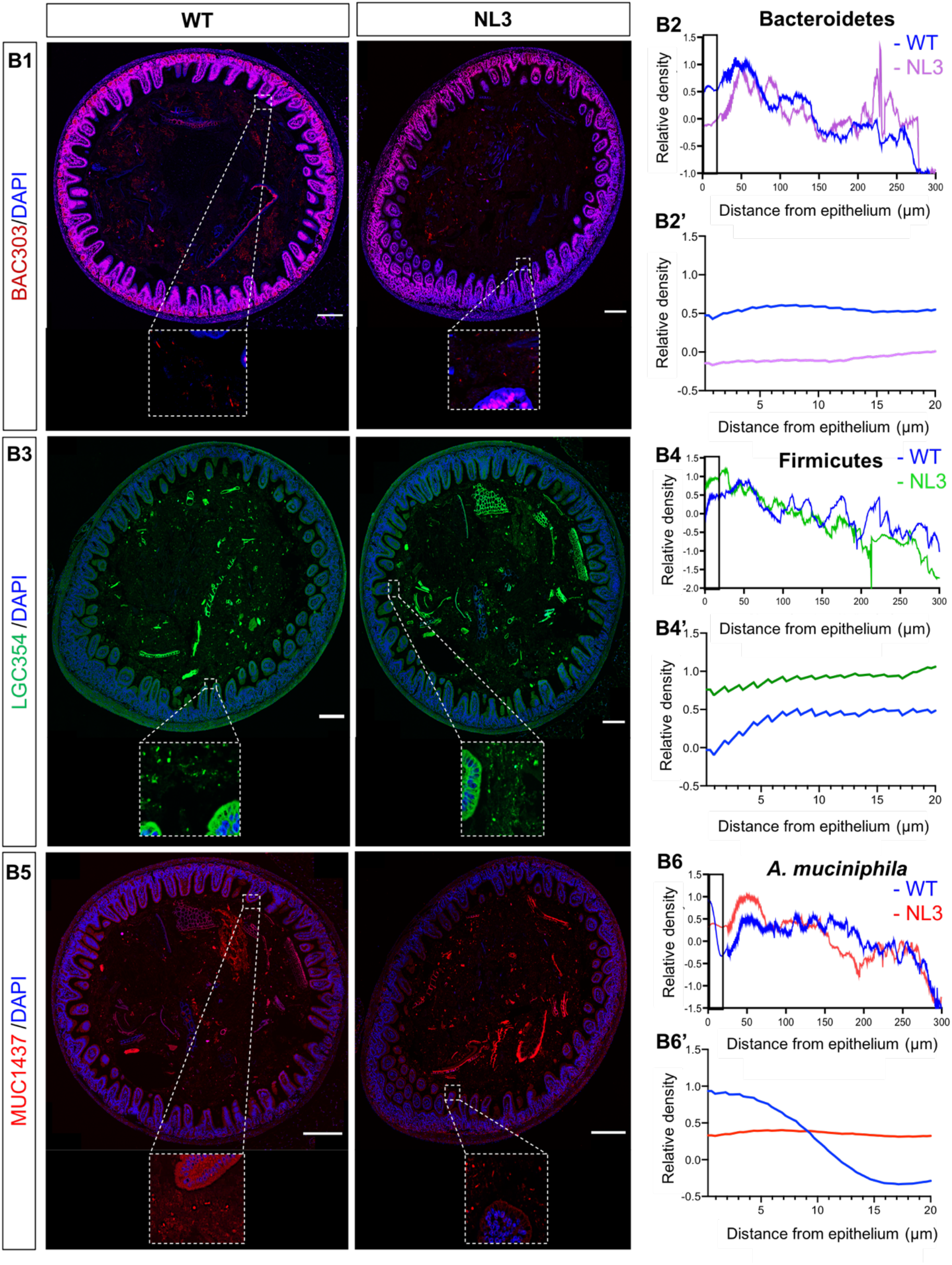
The R451C mutation drives changes in Bacterial localization in the distal ileum. **B1:** Confocal images of distal ileum labelled with the BAC303 probe for the phylum Bacteroidetes and DAPI, a DNA marker for host nuclei in WT and *Nlgn3*^R451C^ mutant mice. White inset: bacterial density adjacent to the epithelium. **B2**: Spatial distribution of Bacteroidetes. **B2’:** (enlarged area of interest indicated by black rectangle in B2): Bacteroidetes density within 20 μm from the epithelium. **B3:** Confocal micrographs of distal ileum samples labelled with LGC354 FISH probe for the phylum Firmicutes and DAPI in WT and *Nlgn3*^R415C^ mice. White insets: bacterial density adjacent to the epithelial boundary. **B4**: Spatial distribution of Firmicutes. **B4’:** (enlarged area of interest indicated by black rectangle in B4): the distribution of Firmicutes organization within 20μm of the epithelial boundary. **B5:** Confocal micrographs of the distal ileum labelled with MUC1437 probe for *A. muciniphila* and DAPI, DNA marker in (**B5**) WT and *Nlgn3*^R451C^ mutant mice. **B6**: Spatial localization of *A. muciniphila* in WT and *Nlgn3*^R451C^ mice. **B6’:** (enlarged area of interest indicated by black rectangle in B6): *A. muciniphila* density from 20 μm adjacent to the epithelium. Data are mean-subtracted and divided by the standard deviation for normalization. Scale bar=100 μm

Although microbial dysbiosis has been identified in people diagnosed with ASD, the spatial localization patterns of bacterial communities within the intestine have not been reported in this population. Dysbiosis in ASD patients has predominantly been identified using 16s rRNA sequencing to measure the microbial abundance in faecal and mucosal samples which does not yield spatial information. The current study showed that the bacterial spatial distribution adjacent to the mucosa in the distal ileum is similar in WT and *Nlgn3*^R451C^ mutant mice when the bacterial density profile from the mucosa extending to 300 microns inside the lumen was examined. Remarkably, however, bacterial density and localization patterns in the region adjacent to the mucosa (i.e., 0-20 microns from the mucosal barrier) were disturbed by the *Nlgn3*^R451C^ mutation. Recently, microbial sequencing studies have generated invaluable information to shed light on how individual bacterial group abundance is altered in the GI tract, specifically in disease states. But, to elucidate dynamic host-microbial interactions, it is essential to determine the location of these bacteria, how their function is connected to the local habitat and how global spatial patterning of these bacteria is altered in disease states. In the intestine, alterations in the mucosal microbiome are highly likely to occur in tandem with changes in mucus layer properties because some mucolytic bacteria use mucus as the sole source of energy and nutrition. The mucus layer analysis indicated that mice expressing the *Nlgn3* R451C mutation have an increased mucus density adjacent to the distal ileal mucosal epithelium. Given that bacterial retention is commonly associated with increased mucus density in patients with small intestinal bacterial overgrowth (Pimentel et al., 2002, Dukowicz et al., 2007) we postulate that mucus accumulation may result in bacterial retention and increased bacterial density adjacent to the mucosa in the *Nlgn3*^R451C^ mouse ileum.

An imbalance in Bacteroidetes and Firmicutes phyla has been previously reported in ASD (Strati et al., 2017, Gibiino et al., 2018). Similar to the current findings in the *Nlgn3*^R451C^ model, several studies reported a lower Bacteroidetes: Firmicutes ratio in ASD populations; whereas other studies showed solely a significant increase in Firmicutes and/or a reduction in Bacteroidetes (Williams et al., 2011, Tomova et al., 2015, Zhang et al., 2018).

Our findings showed that the density of *A. muciniphila* is inversely correlated with mucus density in *Nlgn3*^R451C^ mutant mice. Within the mucus layer, pores in the mucus mesh allow bacteria to penetrate the mucus layer and access various carbohydrates including O-glycan for use as nutrients and as an energy source (Johansson et al., 2011). Some mucolytic species including *A. muciniphila* exclusively feed on mucus O-glycans (Derrien et al., 2004). When the mucus structure is not fully expanded, these bacteria cannot penetrate the mucus mesh, thus limiting their access to mucus-associated energy sources including O-glycans and other carbohydrates. Therefore, a lack of mucus expansion might cause a reduction in the density of mucolytic bacteria associated with the mucosa. Taken together, improper mucus expansion may drive mucus accumulation and the reduction in mucolytic mucus-residing bacteria observed in the distal ileum of *Nlgn3*^R451C^ mice. However, the impact of this mutation on mucus layer structure and composition has not been studied. Therefore, further work is necessary to elucidate biological mechanisms underlying neurally-mediated mucus regulation in ASD. In summary, this study revealed an increase in the microbial density of the mucus layer adjacent to the intestinal epithelium in mice expressing the *Nlgn3* R451C mutation. These findings suggest that mucus accumulation in the mucosa could contribute to mucosal microbial dysbiosis in ASD.

